# Metagenome Assembled Genomes (MAG) Facilitate a Better Understanding of Microbially-mediated Heavy Metal Resistance in Soils from a Former Nuclear Materials Production Facility

**DOI:** 10.1101/2023.10.20.563326

**Authors:** Navya Kommu, Paul Stothard, Christian Chukwujindu, Ashish Pathak, Ashvini Chauhan

## Abstract

Shotgun metagenomes is a repository of all the genes present in an environmental sample. With recent advancements in bioinformatic techniques, it is now possible to in-silico retrieve sequences that belong to specific taxa, followed by assembly and annotation and the obtained sequences are called as metagenome-assembled genome (MAG), which facilitates better understanding of metabolic and other traits without having to culture the microorganism. We applied the MAG technique using the nf-core/mag pipeline on shotgun metagenome sequences obtained from a soil ecosystem that has long-term co-contamination with radionuclides (mainly uranium), heavy metals (mercury, nickel etc.) and organic compounds. Annotation of MAGs was performed using SPAdes and MEGAHIT and genomes were binned and taxonomically classified using the GTDBTk and CAT toolkits within nf-core/mag. Additional annotations were done using Prokka and Prodigal and the dRep program was used to choose specific MAGs for further analysis. Initial analysis resulted in a total of 254 MAGs which met the high-quality standard with the completeness > 95% and contamination < 5%, accounting for 26.67% of all the MAGs (Fig SI-1). After bin refinement and de-replication, 27 MAGs (18 from Winter season and 9 from Summer season) were reconstructed. These 27 MAGs span across 6 bacterial phyla and the most predominant ones were Proteobacteria, Bacteroidetes, and Cyanobacteria regardless of the season. Overall, the *Arthrobacter* MAG was found to be one that was robust for further analysis. Over 1749 genes putatively involved in crucial metabolism of elements viz. nitrogen, phosphorous, sulfur and 598 genes encoding enzymes for metals resistance from cadmium, zinc, chromium, arsenic and copper. In summary, this project enhances our understanding of genes conferring resistance to heavy metals in uranium contaminated soils.

## Introduction

Uranium is a naturally occurring radionuclide associated with volcanic rocks, black shales, or granite at a concentration between 2 and 4 mg/kg of soil, and it can be rendered mobile upon exposure to oxidative groundwater (Nolan & Weber, 2015). The US Environmental Protection Agency sets the maximum allowable uranium contamination level in groundwater to not exceed 30 µg/l; however, anthropogenic activities, including mining, nuclear fuel production, and testing, has historically led to the release of uranium into groundwaters and soils, thus escalating environmental health concerns. Uranium exposure causes pathological alterations in the kidney (nephrotoxic) of both humans and animals (Vincente-Vincente *et al*., 2010), and it accumulates in bone, leading to a widespread osteotoxicity in exposed cohorts, among other serious health issues associated with uranium exposure (Kurttio et al., 2005).

Over 50 uranium companies were involved in uranium processing and refining in the US during the cold war but plans for the treatment of the toxic byproducts were minimal, leading to the seepage of radioactive materials into nearby communities, streams, and aquifers (Olalde et al., 2022). One such former nuclear production facility is the Savana River Site (SRS), covering about 310 square miles, and was built during the 1950s in the upper Atlantic coastal plain of South Carolina. Particularly, Steed Pond, an abandoned farm pond at the Northwest of the SRS, served as a settling basin for contaminants originating from the target production activities and received an estimated 44,000 kg of depleted uranium, leading to high uranium tailing diffusing into the environment, the surface and groundwaters; Punshon et al., 2003 and Sowder et al., 2003, reported that uranium contamination from the Steed Pond sediments could be as high as 2500 mg/kg of soil, about 4th orders of magnitude above the background concentration of 4 mg/kg from the uncontaminated areas. Therefore, investigating approaches to remediate uranium contamination from the SRS soil habitat has been a large focus of our project.

In this context, microbially mediated remediation strategies are cost-effective, eco-friendly, and efficient compared with other remediation strategies (Banala *et al*., 2021). Previous studies have observed relatively high bacterial diversity in samples with different levels of uranium contamination, which suggests that bacterial communities can acclimatize and colonize uranium-contaminated soils and may also exert remediation influences, mainly by immobilizing uranium through adsorption, precipitation, intermolecular interactions, and/or redox transformations (Yan & Luo, 2015; Merroun & Selenska-Pobell, 2008). As part of our broader scale project, we have evaluated bacterial and fungal communities within the SRS contaminated soils using culture-based and culture-independent techniques, which has revealed that

Our ongoing culture-dependent and culture-independent work on the SRS metalliferous soils has demonstrated several different bacterial groups, predominantly from the following genera: *Arthrobacter* spp., *Burkholderia* spp., *Bacillus* spp., *Bradyrhizobium* spp., *Pseudomonas* spp., *Lysinibacillus* spp., *Paenibacillus* spp., *Stenotrophomonas* spp., and *Serratia* spp. (Agarwal et al., 2020a, b; Gendy et al., 2020b; Pathak et al., 2017; Pathak et al., 2020; Jaswal et al., 2019; Chauhan et al., 2017). It is noteworthy that researchers from another DOE site in Washington State (the U-contaminated Hanford site), having a similar uranium contamination history as the SRS site, reported *Arthrobacter* spp., to be predominant (Katsenovich et al., 2013). *Arthrobacter* spp., have also been isolated from ecosystems under extreme environmental stresses, such as a nuclear waste plume and U mined site, as well as heavy metal contaminated habitats. It is very likely that *Arthrobacter* spp., have acquired robust genomic traits to facilitate their survival under heavy metal stress. In this regard, Suzuki and Banfield (2004) showed that *Arthrobacter spp.* accumulate uranium intracellularly in the form of precipitates closely associated with polyphosphate granules, thus performing natural attenuation on sites laden with uranium contaminations. Similarly, Bader et al. (2018) and Banala et al. (2021), showed that *Actinomycetes*, such as *Arthrobacter spp*., have the highest uranium sorption capacity of up to 971 mg U/g among several tested strains. *Actinomycetes* are gram-positive, ubiquitous in soil ecosystems where they perform critical ecological roles, including decomposition of organic materials, lignin, and chitin (Sullivan & Champman, 2015). A better understanding of the basis of metal-microbial interactions can be instrumental towards successful restoration of nuclear-legacy contaminated environments.

The difficulty in culturing *Actinomycetota* and other microbial species that play significant roles in uranium remediation underscores the need to apply Metagenomics Assembled Genome (MAG) techniques to obtain a comprehensive understanding of microorganisms and their environmentally-relevant functions represented in shotgun genome sequence libraries, as the one available through our project from samples collected in summer and winter seasons, from low, medium and high levels of uranium contamination. Furthermore, reference genomes for many uncurable microbes are essential for a detailed functional and taxonomic characterization of species in a microbial population of a given niche, but a vast majority of microbes cannot be isolated for individual sequencing; thus, the MAG technique becomes a viable alternative (Zhou, Liu, & Yang, 2022). Therefore, the main objective of this study was to reconstruct microbial genomes from shotgun metagenome sequence data we have obtained from our ongoing project focused on obtaining a better understanding of heavy metal resistant genes in historically contaminated soils (Pathak et al., 2020; Jaswal et al., 2019 a,b).

## Materials and Methods

### Sample Collection

Samples were collected from the SRS Steeds Pond-Tims Branch riparian stream system, as shown in our previous report (Pathak et al., 2020). Surface soil samples were collected in duplicates from three Steed Pond locations during the winter (January 2016) and the summer (July 2017) season. Samples were stored in sterile containers and transported to FAMU for the microbial analysis.

### Shotgun Metagenomics and Bioinformatics Processing

DNA was extracted from the duplicate samples representing high, medium and low uranium contamination using DNeasy Powersoil Kit (QIAGEN Inc., Germantown, MD, USA) per the manufacturer’s instruction. A NanoDrop 1000 (NanoDrop Technologies, Wilmington, DE, USA) was used to quantify the total DNA of the extracted samples by measuring the concentrations (ng/µL) by absorbance at A260/280, and A260/230 ratios. An aliquot of high-quality genomic DNA was processed for the shotgun sequencing in accordance to protocols described by Illumina Inc. (USA). The equimolar libraries were pooled and sequenced on Illumina HiSeq platform using 2x150bp chemistry.

The processing of the obtained raw reads involved mapping to the NCBI non-redundant (NR) protein database using DIAMOND using default parameters (Buchfink, et al., 2015) followed by taxonomic assignment using MEGAN’s Least Common Ancestor algorithm (Huson, et al., 2007). The raw read counts were transformed to obtain relative abundance which was then subjected to filtration based on the taxonomic rank of kingdom such as bacteria, archaea, viruses, fungi, and non-fungal eukaryotes which yielded respective abundance tables. Functional profiling of these datasets was carried out using SUPER-FOCU to obtain estimated abundance of genes and subsystem pathways which were used to represent persistence of different pathways and their specificity.

### Bioinformatics Analysis to Reconstruct Metagenome Assembled Genomes (MAGs)

The raw shotgun sequence reads were downloaded from the NCBI genome repository (https://www.ncbi.nlm.nih.gov/bioproject/PRJNA436168), as stated below:

1. Soil from summer with high U contamination: SRR6788480, SRR6788481, SRR6788482
2. Soil from summer with medium U contamination: SRR6788478, SRR6788479, SRR6788487;
3. Soil from summer with low U contamination: SRR6788483, SRR6788484, SRR6788485;
4. Soil from winter with high U contamination SRR6788486;
5. Soil from winter with medium U contamination SRR6788488; and
6. Soil from winter with low U contamination SRR6788489

The MAG pipeline nf-core (https://nf-co.re/mag) was ran on the soil data and genome assembly was then done using the programs SPAdes and MEGAHIT. Due to the size, data had to be processed in groups based on sample type. The assembled sequences generated by SPAdes and MEGAHIT were evaluated using the computational tools QUAST. The genome binning was then performed using MetaBAT2, and the quality of the bins, including the completeness and the contamination levels, was evaluated using CheckM and BUSCO. The CheckM analysis also performed the taxonomic classification of the bins.

Once sequence binning of the MAGs was complete, we performed an additional process using the program dRep, which compared the obtained MAGs, enabling us to choose the best representative for each genome. dRep was also utilized to observe which MAGs are closely related and to further analyze genomes of interest. Prokka and Prodigal were used to predict the protein coding genes of the interested MAG. The bins were assigned taxonomic classification using the GTDB-Tk and CAT toolkits. Other annotation workflows such as rapid annotation using subsystems technology (RAST; Aziz et al., 2008), the Pathosystems Resource Integration Center (PATRIC; Wattam et al., 2017) and IslandViewer (Bertelli et al., 2017) were utilized for further genomic analysis on MAGs of specific interest.

## Results and Discussion

### Characteristics of the MAGs recovered from shotgun metagenomes using the CheckM pipeline

Genes assembled using MAGs have lower quality compared to the isolated genomes because MAGs are a mix of genes from different microbial populations, but this technique has contributed to the discovery of a vast array of novel organisms and functions from diverse samples, including guts, rumens, sediments, soil, with a three-step basic conventional workflow: preprocessing of sequence reads, construction of MAGs, and quantification and annotation of MAGs (Saheb Kashaf et al.,2021; Zhou et al., 2022). Using a MAG approach, a total of 55681 Mbytes of sequences were obtained from 12 soil samples, having a variable level of uranium contamination. Binning of the metagenomics assembled sequences was carried out with MetaBAT pipeline and the quality assessment of the obtained bins was by CheckM, all within the nf-core domain. CheckM does a phylogenetic placement of the bin genomes into its separate species tree which allows the computation of the universal and additional single copy marker genes specific for a particular lineage, giving rise to the completeness of the bins (Parks et al., 2014). About 254 individual bins were recovered from the SRS soil samples; however, due to the sheer volume of data retrieved, the data table was limited to MAGs produced with >50% completeness (Fig. SI-1), that possessed unique marker lineages representative of the range of genomes present in the samples. As shown in table SI-1A, the assembled sequences generated by SPAdes and MEGAHIT were evaluated using the the <u>QU</u>ality <u>AS</u>sessment <u>T</u>ool (QUAST) computational pipeline. The genome binning was then performed using MetaBAT2, and the quality of the bins, including the completeness and the contamination levels, was evaluated using CheckM and BUSCO, thus estimating the completeness and redundancy of processed genomic data based on universal single-copy orthologs (table SI-1B).

Furthermore, as shown in table 1, the CheckM analysis represented important features of metagenome assembled genomes (MAGs) reconstructed from the SRS soil samples. Specifically, the quality assessment with CheckM reduced the number of bins to 25, distributed at 4, 4, 4, 5, and 8 for the summer high, summer low, winter high, winter low, and winter medium uranium-contaminated soil samples, respectively. Moreover, out of the 25 bins that passed the completeness threshold, 5 were classified to the phylum level, 4 to the class level, one to the order level, and the rest were at the kingdom level, suggesting that the MAG technique for the assessment of the SRS soil samples likely captured un-culturable bacteria not represented in the database. In addition, two of the 5 phyla belonged to *Actinobacteria,* which have been previously identified from our previously published findings on the uranium contaminated SRS ecosystem (Chauhan et al., 2018; Jaswal et al., 2019), thus the MAG further highlights the potential role of this phyla in heavy metal resistance processes.

At a broader scale, phylogenomic analysis based on sets of single copy marker genes universal to the bacterial or archaeal domains showed that the 254 MAGs reconstructed from SRS soils regardless of seasons or contamination levels, were distributed in the following taxonomic groups: *Alphaproteobacteria, Gammaproteobacteria, Deltaproteobacteria, Betaproteobacteria, Euryarchaeota, Archaea, Acidobacteria, Actinobacteria, and Rhizobiales*. These findings are in line with our previous reports on the taxonomic assessment of the SRS soils (Pathak et al., 2020; Jaswal et al., 2019 a,b).

### Statistical Analysis of the MAGs from SRS Uranium Contaminated Soils

The recovered MAGs were analyzed statistically using MASH to evaluate which MAGs are closely related and were plotted using a dendrogram (Figure 1). MASH tool extends the MinHash dimensionality-reduction technique to include a pairwise mutation and p-value significance test for efficient clustering of sequence collections (Ondov et al., 2016). However, the dendrogram analysis revealed that the MAGs did not cluster based on the uranium concentration or the sampling season. This dendrogram was constructed after the program dRep was used to identify sets of genomes that are similar and choosing the best representative genome from each set (referred to as the “winning genome”). dRep is a metagenomics tool that identifies groups of essentially identical genomes from independently assembled genomes of metagenomic samples and selects the best genome from each replicate set (Olm et al., 2017). dRep algorithm combines different metagenomics tools, such as CheckM for completeness-based genome filtering, Mash for fast grouping of similar genomes, gANI for accurate genome comparison, and Scipy for hierarchical clustering (Olm et al., 2017). The 254 MAGS from SRS shotgun metagenomes were compared and processed to choose all the winning genomes. Analysis of the winning genomes revealed the predominance of *Arthrobacter oryzae,* thus pointing to the relevance of this soil borne-bacterial genera in the SRS historically contaminated soils.

**Fig. 1.**
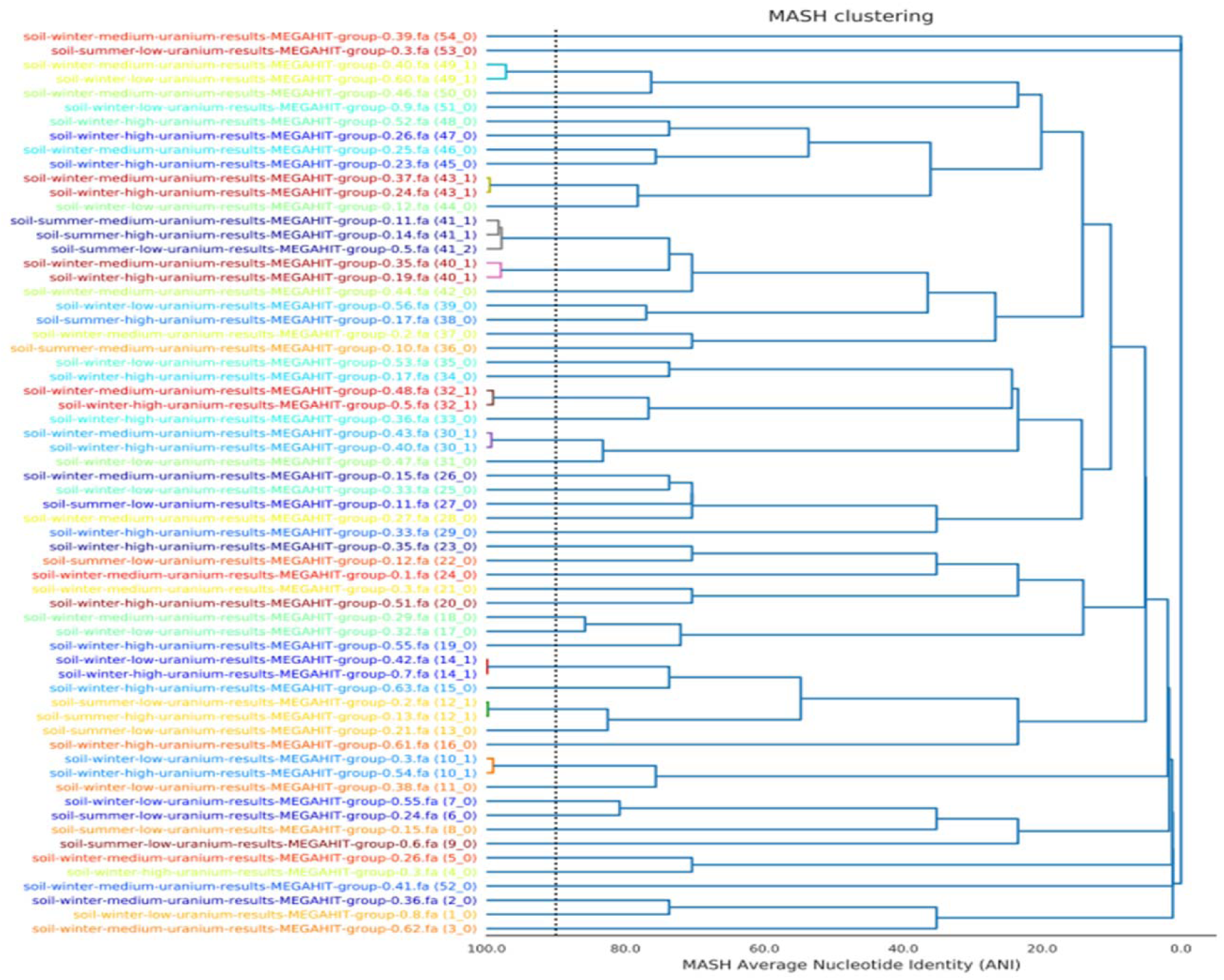
Dendrogram of the best representative MAGs from SRS uranium-contaminated soil samples. The dendrogram was constructed using MASH based on the average nucleotide identity (ANI) to illustrate which MAGs are closely related. The dendrogram revealed that the MAGs did not cluster based on uranium concentration or the sampling season.

### Assessment of Metabolic Traits of the Winning MAG -*Arthrobacter oryzae*

RAST is a fully automated service for annotating complete or nearly complete bacterial or archaeal genomes by identifying protein-encoding, rRNA and tRNA genes, assigning functions to the genes and predicting which subsystems are represented in the genomes (Aziz et al., 2008). Figure 2 shows a metabolic model constructed from the SRS MAG belonging to *Arthrobacter oryzae*. This MAG was specifically chosen for further genomic analysis as *Arthrobacter* is one bacterial genus previously identified to perform critical role(s) in heavy metal resistance within the SRS soil habitat (Chauhan et al., 2016). This model obtained using the RAST in-silico workflow (Surachat et al., 2022) revealed that only 26% of the genes could be assigned to known functions and 74% were found to not have any affiliation with biologically known functions. This indicates that contaminated soil environments could potentially be driving the evolution of microbes with genomic traits that are hitherto unrepresented in the annotation databases.

**Fig. 2.**
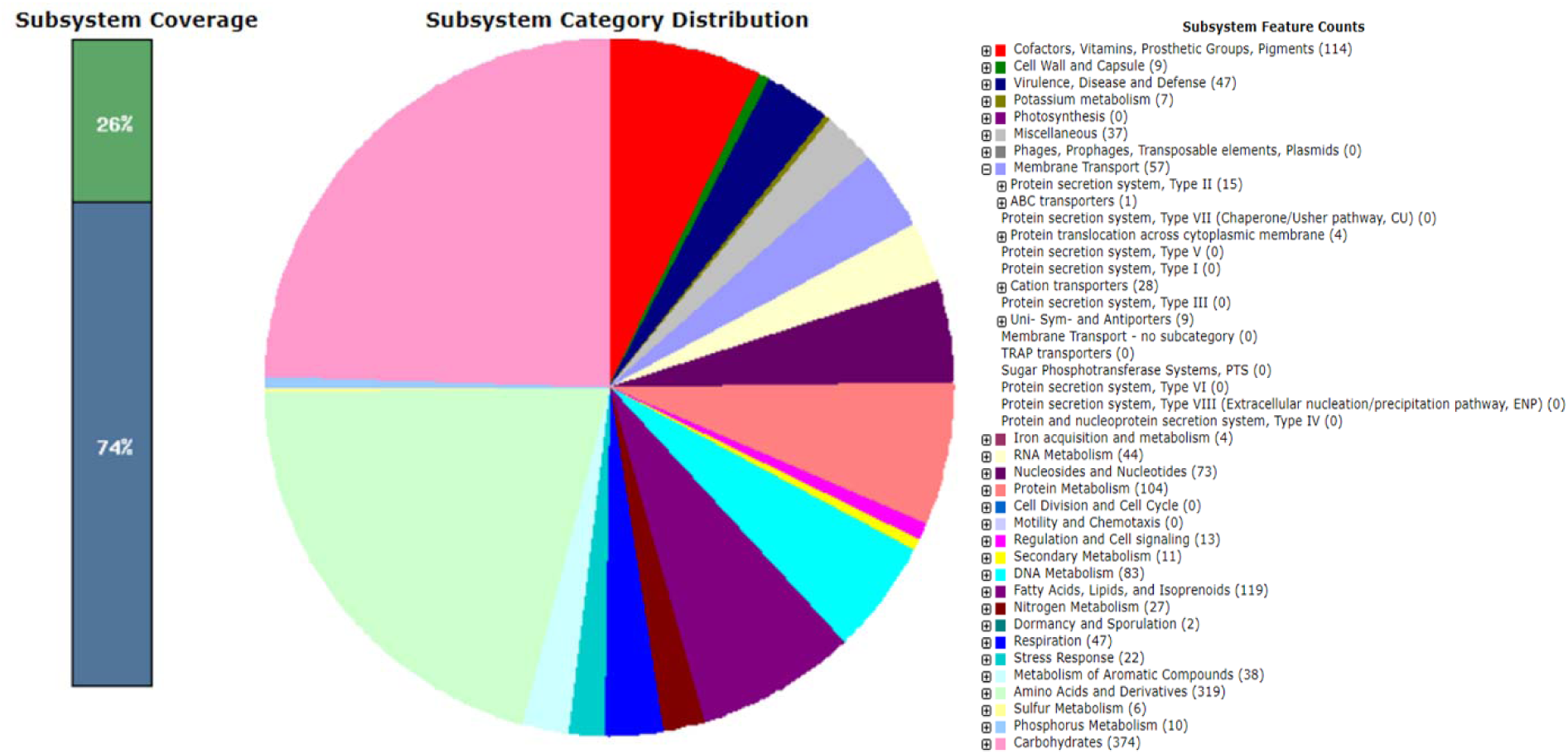
Metabolic traits of the MAG (*Arthrobacter oryzae)* from SRS uranium contaminated soil. The Metabolic trait of Arthrobacter oryzae was computed using Rapid Annotations using Subsystems Technology (RAST). The model chart indicates that 26% of the genes could be assigned to known functions with 74% with no affiliation to known functions. Carbohydrate metabolism, amino acid and derivatives, protein metabolism, cofactor and vitamin metabolism, and membrane transport are among the major gene classes identified in this species.

Most notably, carbohydrate metabolism, protein metabolism, cofactor and vitamin metabolism, and membrane transport were among the major gene classes identified in the MAG (Figure 2). These features suggest the tendency of *Arthrobacter oryzae* to participate in high metabolic and catabolic activity in its contaminated habitat which likely plays a key role in its survival and ability to degrade harmful pollutants. Membrane transport is especially important for bioremediation because it allows microorganisms to regulate their internal environment by extruding toxic chemicals, such as uranium, thereby maintaining cytoplasmic contents and homeostatic balance necessary for growth (Guan, et al., 2022).

### Gene Functional analysis of the *Arthrobacter oryzae* MAG

Gene functional analysis was performed on the *Arthrobacter oryzae* MAG, and is shown in Figure 3, obtained using the One Codex bioinformatics pipeline. It is clear from this analysis that the ATP-binding cassette family (ABC), as well as the Major Facilitator Superfamily (MFS) are among the two most overrepresented genes in the SRS *Arthrobacter* MAG. ABC transporter proteins and the MFS are two of the five major families of efflux transporters (ref), are thus likely important for rendering heavy metal resistance to the native SRS soil microbiota. According to Akhtar and Turner (2022) ABC transporters superfamily are transporters found in all domains of life, translocating various molecules, such as toxins and xenobiotics, to the outside of the cell’s cytoplasm, or they can even import various nutrients for cellular survival. In the same way, the Major Facilitator superfamily (MFS) is another membrane transporter that plays a key role in the maintenance of cellular elements. Pao et al (1998) reported the occurrence of several distinct families within the MFS, each of which generally transporting a single class of organic and inorganic anions and cations. Note that our previous analysis on heavy metal contaminated soils have also shown the ABC and MFS to be predominant in historically contaminated soils. Moreover, the analysis performed on the SRS *Arthrobacter* MAG using the KBase bioinformatics pipeline yielded an abundance of genes notable to DNA recombination and repair, as well as defense and virulence responses, as shown in table SI-2. A plethora of defense and repair mechanisms is essential to counter the damage this organism may face from constant exposure to a heavy metal contaminated environment. Furthermore, this species was shown to contain several metal transporting ATPases and metal efflux proteins, that are important adaptations to survive in a heavy metal contaminated environment and enhance the heavy metal resistance systems this organism harbors (Hemme et al, 2010). If the metabolic efficacy of the above stated gene classes are enhanced in contaminated environments, perhaps bioremediation of heavy metals can be rapidly accomplished for better management of contaminated habitats.

**Fig. 3.**
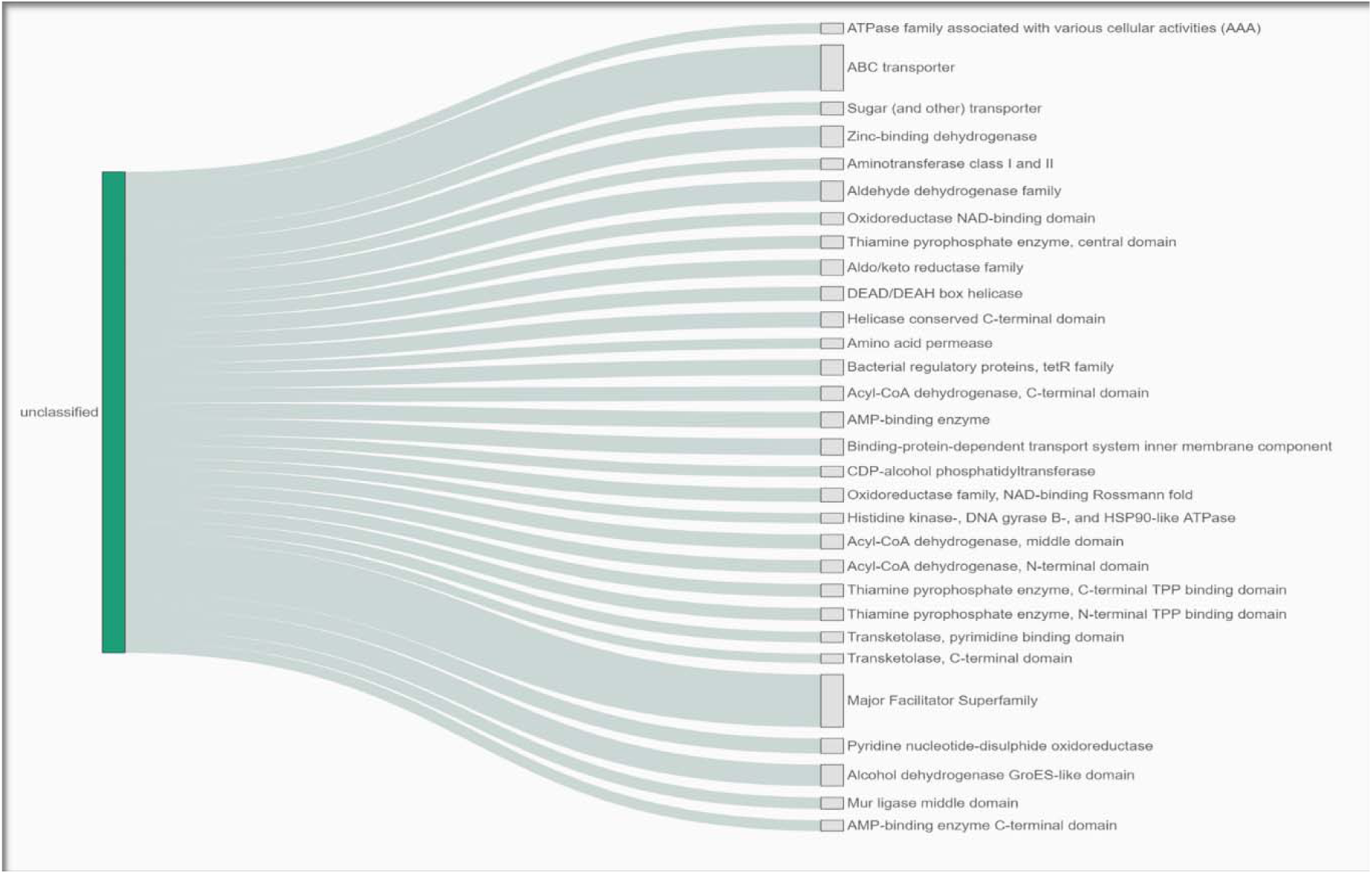
Functional analysis of the *Arthrobacter oryzae* genome to examine specific genes that aid in uranium bioremediation. The uranium bioremediation functional analysis was carried out using One Codex bioinformatics pipeline. The ATP-binding cassette family (ABC), and the Major Facilitator Superfamily are among the most overrepresented genes in this species. ABC transporter proteins and the Major Facilitator Superfamily are two of the five major families of efflux transporters. Efflux transporters are important for bioremediation because they can, not only prevent contaminants from entering a cell, but also convert harmful compounds within a cell to a more hydrophilic form.

### Genome Map of *Arthrobacter oryzae MAG* Reconstructed from the SRS Uranium Contaminated Soil

Figure 4 shows the circular genome map of *Arthrobacter oryzae* MAG depicting DNA contigs, AMR Genes, VF Genes, Drug Targets, GC Content, GC Skew, Protein families, Efflux pumps, Permease proteins, and Transporters, obtained using the Pathosystems Resource Integration Center (PATRIC) pipeline. PATRIC is a bacterial bioinformatics pipeline that can be used to run a variety of genomic analysis including genomic assembly using different strategies, genomic annotation using RAST tool kit, and provision of several annotated genomes for comparative studies (Wattam et al., 2018). The Circular Viewer tool in PATRIC provides the circular genomic map using all the contigs from an assembly via the Circos technology (www.patricbrc.org). The genomic map shown in Figure 4 consists of 13 circles from outside in; the first shows size in Mbp, followed by DNA contigs, AMR Genes, VF Genes, Transporters, Drug Targets, GC Content, GC Skew, Protein families, Efflux pumps, Permease proteins, and Transporters.

**Fig 4.**
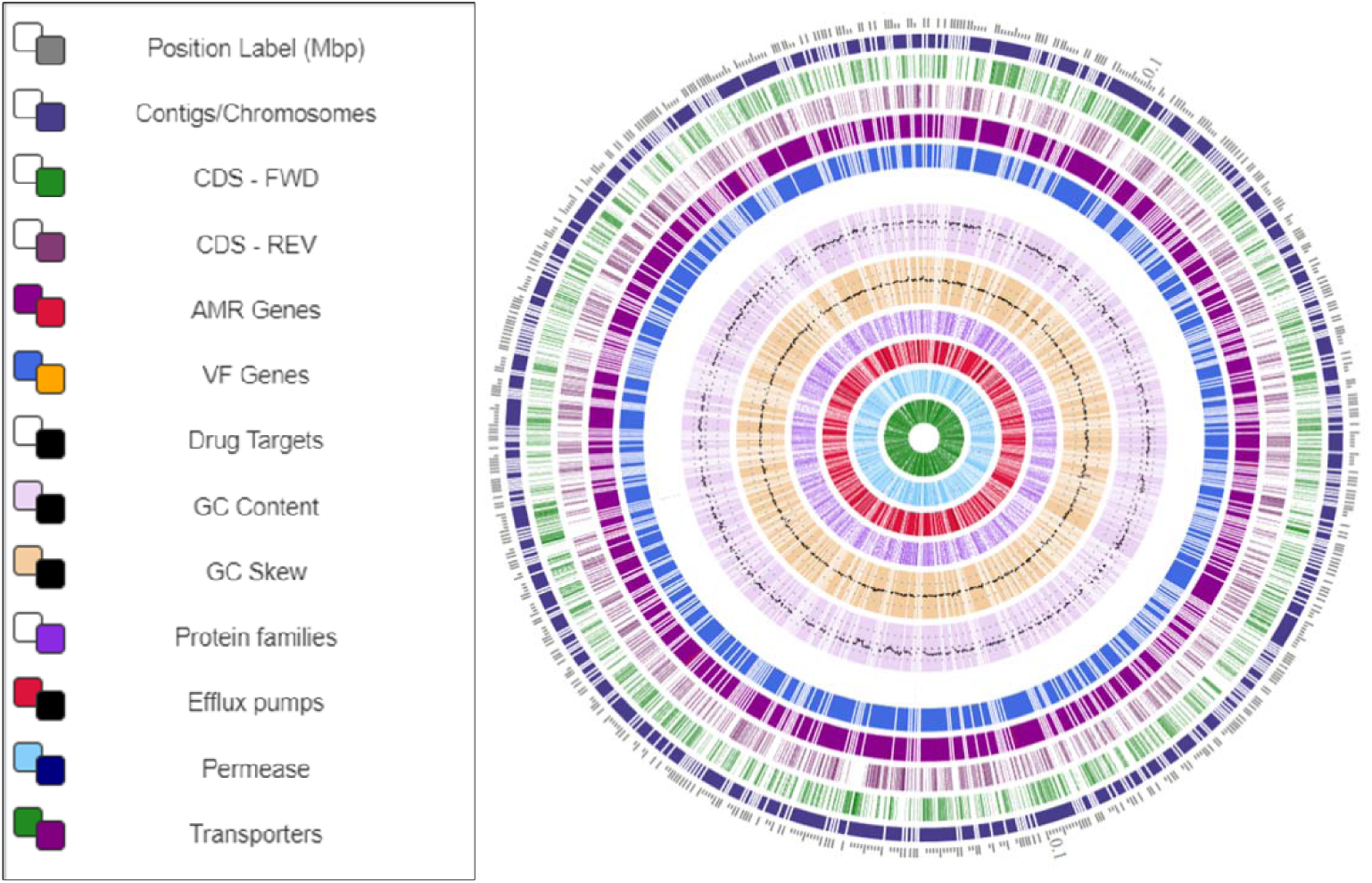
Circular genome map using PATRIC default settings to visualize the genomic features of *Arthrobacter oryzae* species from SRS uranium contaminated soil. The map consists of 13 circles from outside in; the first shows size in Mbp, followed by DNA contigs, CDS forward and reverse, AMR Genes, VF Genes, Drug Targets, GC Contents, GC Skew, Protein families, Efflux pumps, Permease proteins, and Transporters, respectively.

In the genomic map of the MAG, of particular interest was the identification of several protein families, efflux pumps, and permease proteins, as shown by the white/purple, red/black, and light/dark blue respectively. As stated above, efflux pumps are transport proteins involved in the extrusion of hazardous substrates from within the cells into the external environment, and they are linked to bioremediation mechanisms (Sharma et al., 2023). In addition, permease proteins are essential membrane transport mechanisms that allow the introduction of stronger proteins into the bacterial cell through genetic engineering to enhance bioremediation traits (Rafeeq et al. 2023). Interestingly, Cho et al. (2019) drafted the whole genome of an *Arthrobacter oryzae* strain from a site not affected by anthropogenic activities and reported that the genome contains many heavy metal resistant genes, so bioremediation may be a central genome-enabled trait of this genera.

Moreover, this study also identified several antimicrobial resistance (AMR) genes, depicted by the purple/red lines in the genomic map. Specifically, the number and types of specialized genes found in the MAG of *Arthrobacter oryzae* are shown in the supplementary Table S1-3, thus further supporting our observation that the SRS contaminated site is likely “priming” the evolution of antibiotic resistant microbes; this contributes to the public health threat these contaminated sites pose.

### Genomic Islands (GEIs) in the *Arthrobacter oryzae* MAG

Genomic Islands (GEIs) are clusters of genes in a microbial genome acquired by horizontal gene transfer, carrying genes that are important for genome evolution and adaptation to niches, such as genes involved in heavy metal resistance (Lu & Leong, 2016). Figure 5 shows the identification of several genomic islands (GEIs) in the MAG of *Arthrobacter oryzae*. The outer green circle represents the scale line in Mbps and GEIs obtained from each of the following methods are shown in color: SIGI-HMM (orange), IslandPath-DIMOB (blue), and integrated detection (red). GEIs prediction methods usually exploit sequence composition and sporadic phylogenetic distribution as the main features of GEI in a genome; based on these features, the GEI methods can be either composition based or comparative genomic based (Langille et al., 2010). The SIGI-HMM method uses the Hidden Markov Models to predict the GEIs (Waack et al., 2006), while the IslandPath-DIMOB and integrated detection methods are based on the GEI structure of the genome (Lu & Leong, 2016). The visual in Figure 5 shows several potential genomic islands within the *Arthrobacter* MAG and further analysis of the GEI regions using BLAST showed gene homologs with functions of metal resistance and biodegradation of contaminants. Molecular transport mechanisms, such as ABC transporters and MFS genes as mentioned earlier, can facilitate metabolic processes that transform toxic metal compounds into less detrimental forms, as well as the development of stronger cell walls and membranes used for protection against the toxic effects of heavy metals. These traits can be acquired via horizontal gene transfer processes (Li et al, 2023). Moreover, it has been shown that heavy metal resistance mechanisms can greatly differ between different taxa, even those within the same community (Lin et al, 2021). Thus, it is possible that this *Arthrobacter oryzae* species acquired taxa specific metal resistance traits to allow it to survive in its stressful environment through horizontal gene transfer mechanisms. This suggests acquiring GEIs likely facilitate the survival of microorganisms in a heavy metal and aromatic compound contaminated environment, such as the Savannah River Site (SRS); these findings mirror our previous genome-enabled studies on isolated organisms, such as *Arthrobacter*, *Serratia*, *Stenotrophomonas* and *Burkholderia* from the SRS site (Agarwal et al., 2018; Chauhan et al., 2016; Jaswal et al., 2019 a, b; Pathak et al., 2017; Pathak et al., 2018; Pathak et al). In summation, this MAG based study lends support to our previous studies conducted on the SRS heavy metal resistant microbiota and further enhances our understanding of genes conferring resistance to heavy metals in uranium contaminated soils. These findings can be applied to develop innovative methods to restore heavy metal contaminated sites such as the SRS.

**Fig. 5.**
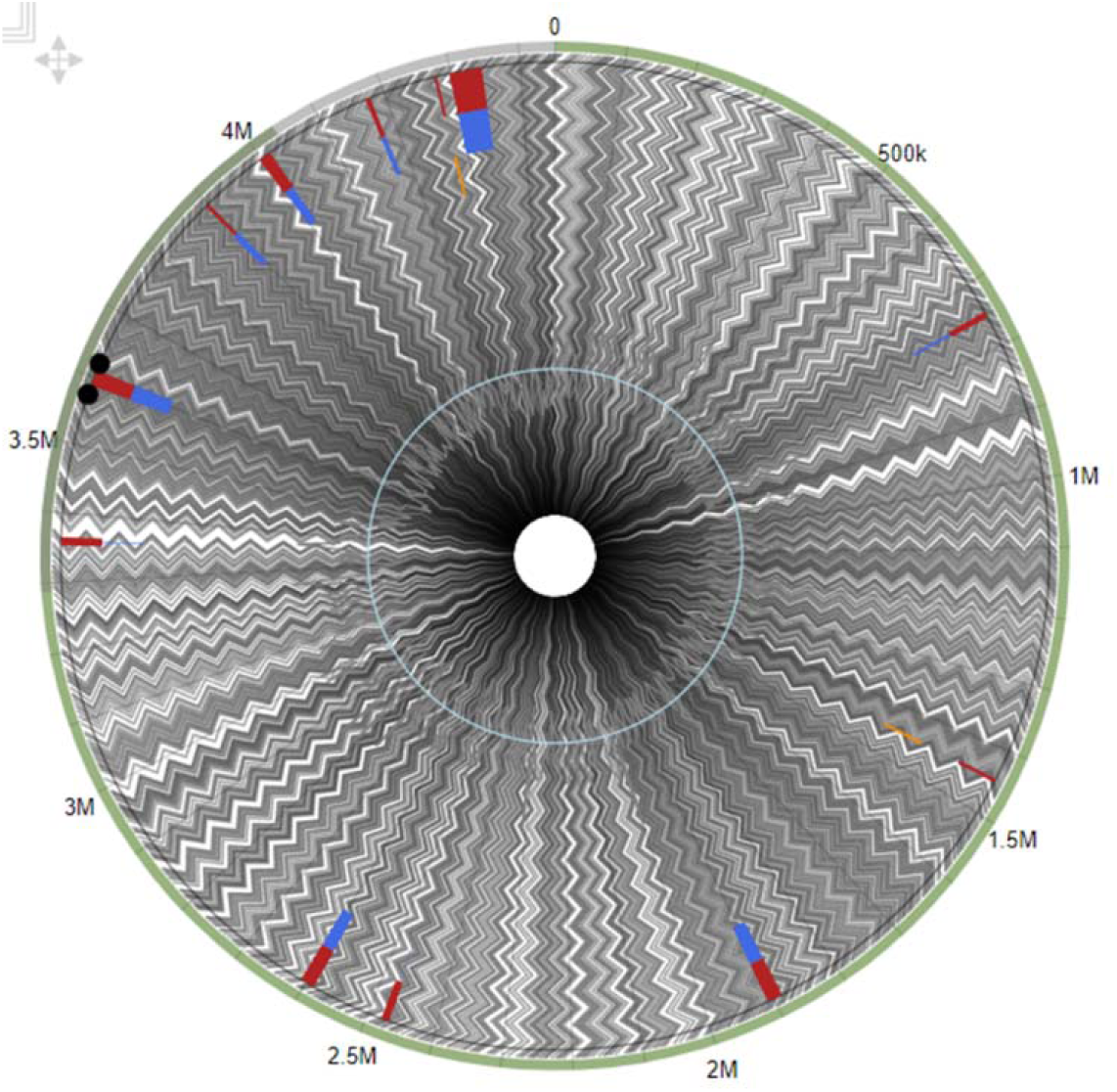
The Genomic Island (GEI) view of *Arthrobacter oryzae* MAG obtained with Islandviewer 4. GEIs obtained from each of the following methods are shown in color: SIGI-HMM (orange), IslandPath-DIMOB (blue), and integrated detection (red), respectively.

---

**Fig. SI-1.**
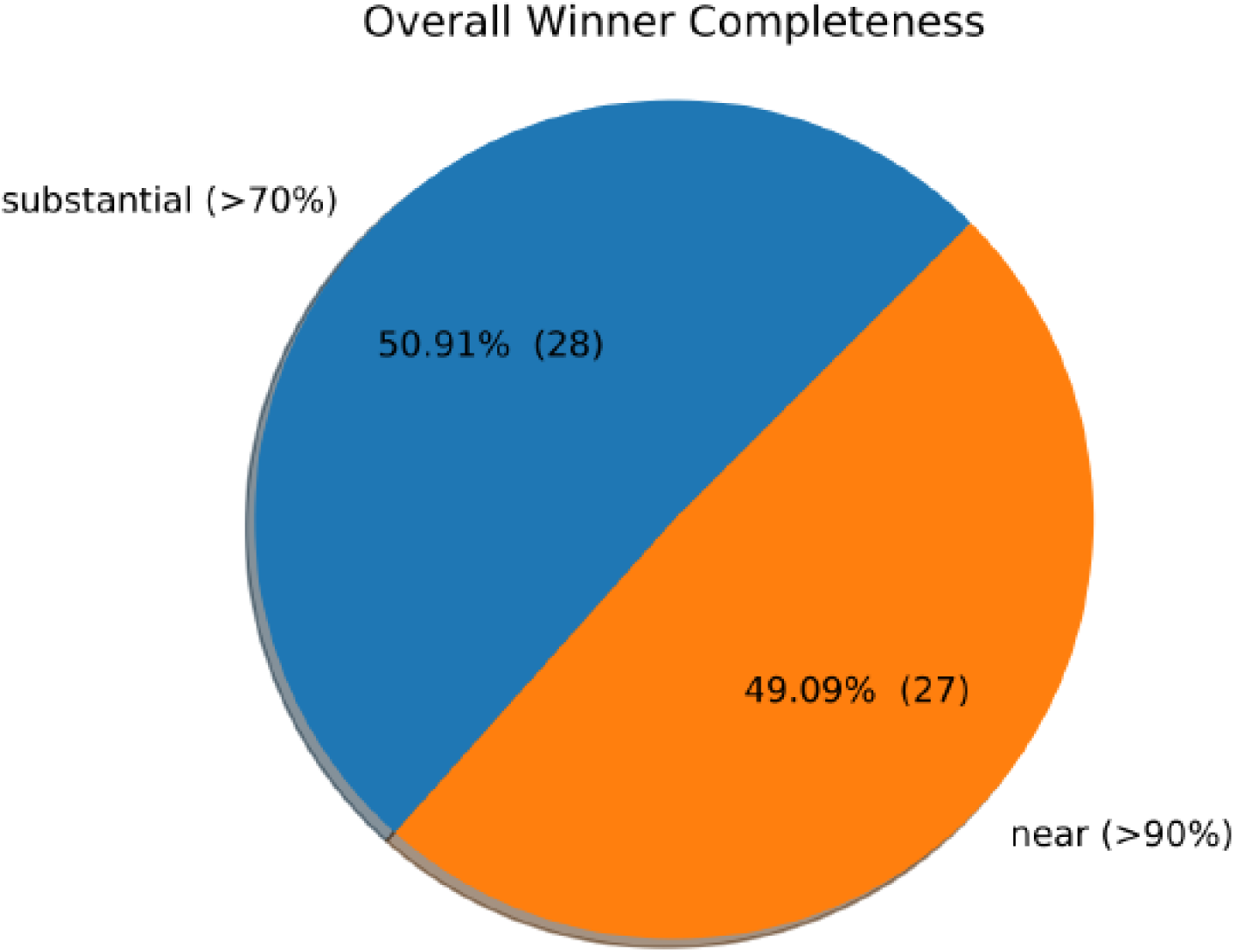
Overall winner MAGs produced from the SRS soils reported in this study.

**Table SI-1A.** Genome summary assessment of MAGs using the <u>QU</u>ality <u>AS</u>sessment <u>T</u>ool (QUAST) pipeline.

| **GenomeBin** | **Specific lineage dataset** | **%Complete (specific)** | **%Complete and single-copy (specific)** | **%Complete and duplicated (specific)** | **%Fragmented (specific)** | **%Missing (specific)** | **Total number (specific)** |
| --- | --- | --- | --- | --- | --- | --- | --- |
| SPAdes-group-0.1.fa | bacteria_odb10 | 30.7 | 22.6 | 8.1 | 10.5 | 58.8 | 124 |
| MEGAHIT-group-0.4.fa | bacteria_odb10 | 0.8 | 0.8 | 0 | 3.2 | 96 | 124 |
| MEGAHIT-group-0.1.fa | bacteria_odb10 | 42.7 | 39.5 | 3.2 | 12.9 | 44.4 | 124 |
| MEGAHIT-group-0.2.fa | bacteria_odb10 | 58.1 | 58.1 | 0 | 11.3 | 30.6 | 124 |
| SPAdes-group-0.11.fa | bacteria_odb10 | 70.2 | 70.2 | 0 | 9.7 | 20.1 | 124 |
| SPAdes-group-0.4.fa | bacteria_odb10 | 30.6 | 30.6 | 0 | 8.1 | 61.3 | 124 |
| SPAdes-group-0.2.fa | bacteria_odb10 | 0 | 0 | 0 | 0.8 | 99.2 | 124 |
| SPAdes-group-0.7.fa | bacteria_odb10 | 76.6 | 75.8 | 0.8 | 0.8 | 22.6 | 124 |
| SPAdes-group-0.6.fa | bacteria_odb10 | 0 | 0 | 0 | 0.8 | 99.2 | 124 |
| MEGAHIT-group-0.5.fa | bacteria_odb10 | 38.7 | 38.7 | 0 | 8.9 | 52.4 | 124 |
| MEGAHIT-group-0.7.fa | bacteria_odb10 | 9.7 | 9.7 | 0 | 5.6 | 84.7 | 124 |
| MEGAHIT-group-0.6.fa | bacteria_odb10 | 0.8 | 0.8 | 0 | 0 | 99.2 | 124 |
| MEGAHIT-group-0.8.fa | bacteria_odb10 | 57.2 | 52.4 | 4.8 | 13.7 | 29.1 | 124 |
| SPAdes-group-0.10.fa | bacteria_odb10 | 0 | 0 | 0 | 1.6 | 98.4 | 124 |
| SPAdes-group-0.8.fa | bacteria_odb10 | 16.1 | 16.1 | 0 | 8.9 | 75 | 124 |
| SPAdes-group-0.9.fa | bacteria_odb10 | 0.8 | 0.8 | 0 | 0 | 99.2 | 124 |
| MEGAHIT-group-0.3.fa | bacteria_odb10 | 79 | 79 | 0 | 2.4 | 18.6 | 124 |
| MEGAHIT-group-0.9.fa | bacteria_odb10 | 4 | 4 | 0 | 4.8 | 91.2 | 124 |
| SPAdes-group-0.3.fa | bacteria_odb10 | 6.5 | 6.5 | 0 | 0.8 | 92.7 | 124 |
| SPAdes-group-0.5.fa | bacteria_odb10 | 38.7 | 38.7 | 0 | 8.9 | 52.4 | 124 |

**Table SI-1B.** Assessment of MAGs using the BUSCO analysis, which estimates the completeness and redundancy of processed genomic data based on universal single-copy orthologs.

| Assembly | # contigs<br>(>= 0 bp) | # contigs<br>(>= 1000<br>bp) | # contigs<br>(>= 5000<br>bp) | # contigs<br>(>= 10000<br>bp) | # contigs<br>(>= 25000<br>bp) | # contigs<br>(>= 50000<br>bp) | Total<br>length<br>(>= 0<br>bp) | Total<br>length<br>(>= 1000<br>bp) | Total<br>length<br>(>= 5000<br>bp) | Total<br>length<br>(>= 10000<br>bp) | Total<br>length<br>(>= 25000<br>bp) | Total<br>length<br>(>= 50000<br>bp) | #<br>contigs | Largest<br>contig | Total<br>length | GC<br>(%) | N50 | N75 | L50 | L75 | # N's<br>per 100<br>kbp | # predicted<br>rRNA<br>genes |
| --- | --- | --- | --- | --- | --- | --- | --- | --- | --- | --- | --- | --- | --- | --- | --- | --- | --- | --- | --- | --- | --- | --- |
| MEGAHIT-group-0.1.fa | 1164 | 1164 | 111 | 10 | 1 | 0 | 3397323 | 3397323 | 828723 | 157449 | 43073 | 0 | 1164 | 43073 | 3397323 | 44.26 | 3044 | 2080 | 344 | 685 | 0 | 1 + 0 part |
| MEGAHIT-group-0.2.fa | 413 | 413 | 30 | 2 | 0 | 0 | 1188763 | 1188763 | 210347 | 24783 | 0 | 0 | 413 | 13354 | 1188763 | 37.78 | 2989 | 2186 | 132 | 250 | 0 | 1 + 2 part |
| MEGAHIT-group-0.3.fa | 497 | 497 | 260 | 140 | 19 | 0 | 4039171 | 4039171 | 3360522 | 2466971 | 593792 | 0 | 497 | 44326 | 4039171 | 40.21 | 12646 | 7181 | 102 | 206 | 0 | 0 + 3 part |
| MEGAHIT-group-0.4.fa | 578 | 578 | 42 | 6 | 0 | 0 | 1524729 | 1524729 | 303252 | 82585 | 0 | 0 | 578 | 19366 | 1524729 | 37.38 | 2589 | 1923 | 179 | 352 | 0 | 5 + 2 part |
| MEGAHIT-group-0.5.fa | 639 | 639 | 29 | 3 | 0 | 0 | 1765814 | 1765814 | 213714 | 49725 | 0 | 0 | 639 | 17739 | 1765814 | 47.62 | 2863 | 2095 | 212 | 393 | 0 | 0 + 0 part |
| MEGAHIT-group-0.6.fa | 24 | 24 | 12 | 6 | 1 | 1 | 220391 | 220391 | 192841 | 151050 | 60563 | 60563 | 24 | 60563 | 220391 | 32.13 | 18239 | 8736 | 4 | 8 | 0 | 0 + 0 part |
| MEGAHIT-group-0.7.fa | 307 | 307 | 1 | 0 | 0 | 0 | 661811 | 661811 | 6111 | 0 | 0 | 0 | 307 | 6111 | 661811 | 42.74 | 2169 | 1773 | 121 | 206 | 0 | 0 + 0 part |
| MEGAHIT-group-0.8.fa | 1242 | 1242 | 169 | 21 | 0 | 0 | 4048292 | 4048292 | 1253825 | 291125 | 0 | 0 | 1242 | 20472 | 4048292 | 42.24 | 3548 | 2390 | 353 | 707 | 0 | 2 + 1 part |
| MEGAHIT-group-0.9.fa | 267 | 267 | 20 | 6 | 0 | 0 | 764945 | 764945 | 165249 | 78021 | 0 | 0 | 267 | 17209 | 764945 | 37.32 | 3017 | 2071 | 77 | 157 | 0 | 3 + 0 part |
| SPAdes-group-0.1.fa | 893 | 893 | 131 | 21 | 0 | 0 | 3004744 | 3004744 | 1029513 | 286339 | 0 | 0 | 893 | 22088 | 3004744 | 44.54 | 3602 | 2390 | 242 | 501 | 263.92 | 1 + 0 part |
| SPAdes-group-0.10.fa | 192 | 192 | 14 | 0 | 0 | 0 | 524106 | 524106 | 94908 | 0 | 0 | 0 | 192 | 8863 | 524106 | 37.35 | 2814 | 2018 | 62 | 116 | 20.99 | 0 + 0 part |
| SPAdes-group-0.11.fa | 317 | 317 | 97 | 20 | 1 | 0 | 1427112 | 1427112 | 792730 | 256455 | 25053 | 0 | 317 | 25053 | 1427112 | 37.82 | 5734 | 3285 | 83 | 164 | 344.75 | 2 + 1 part |
| SPAdes-group-0.2.fa | 136 | 136 | 19 | 4 | 0 | 0 | 435974 | 435974 | 154332 | 56053 | 0 | 0 | 136 | 17152 | 435974 | 37.93 | 3455 | 2160 | 35 | 76 | 25.23 | 0 + 0 part |
| SPAdes-group-0.3.fa | 103 | 103 | 9 | 0 | 0 | 0 | 307428 | 307428 | 59434 | 0 | 0 | 0 | 103 | 8648 | 307428 | 42.42 | 3193 | 2311 | 34 | 62 | 292.75 | 0 + 0 part |
| SPAdes-group-0.4.fa | 474 | 474 | 57 | 7 | 0 | 0 | 1479441 | 1479441 | 407532 | 94802 | 0 | 0 | 474 | 20550 | 1479441 | 43.63 | 3391 | 2228 | 140 | 276 | 229.82 | 0 + 0 part |
| SPAdes-group-0.5.fa | 455 | 455 | 109 | 23 | 0 | 0 | 1873903 | 1873903 | 870720 | 292337 | 0 | 0 | 455 | 17252 | 1873903 | 48.48 | 4616 | 3048 | 123 | 247 | 277.5 | 0 + 0 part |
| SPAdes-group-0.6.fa | 27 | 27 | 17 | 6 | 1 | 0 | 212098 | 212098 | 186230 | 109587 | 29318 | 0 | 27 | 29318 | 212098 | 30.54 | 10507 | 6446 | 6 | 12 | 391.33 | 0 + 0 part |
| SPAdes-group-0.7.fa | 745 | 745 | 267 | 122 | 20 | 1 | 4536590 | 4536590 | 3322404 | 2275821 | 679877 | 56384 | 745 | 56384 | 4536590 | 39.4 | 10014 | 4541 | 122 | 284 | 83.98 | 0 + 0 part |
| SPAdes-group-0.8.fa | 665 | 665 | 31 | 2 | 0 | 0 | 1695251 | 1695251 | 205065 | 25680 | 0 | 0 | 665 | 15228 | 1695251 | 42.42 | 2504 | 1948 | 226 | 418 | 88.48 | 0 + 0 part |
| SPAdes-group-0.9.fa | 24 | 24 | 7 | 5 | 2 | 1 | 221808 | 221808 | 182395 | 167127 | 116746 | 77789 | 24 | 77789 | 221808 | 32.54 | 38957 | 15898 | 2 | 5 | 0 | 0 + 0 part |

**Table SI-2.** Analysis performed on the MAGs of *Arthrobacter oryzae* using the KBase bioinformatics pipeline.

| **Function Role** | **Gene count** | **Subsystem Name** | **Subsystem Class** |
| --- | --- | --- | --- |
| ATP-dependent DNA helicase RecQ | 6 | DNA repair, bacterial RecFOR pathway | DNA processing |
| ATP-dependent DNA helicase SCO5183 | 2 | DNA repair, bacterial RecBCD pathway | DNA processing |
| ATP-dependent DNA helicase SCO5184 | 4 | DNA repair, bacterial RecBCD pathway | DNA processing |
| DNA integrity scanning protein DisA | 2 | DNA repair, bacterial | DNA processing |
| DNA polymerase like protein MT3142 | 4 | DNA repair, bacterial | DNA processing |
| DNA recombination and repair protein RecF | 2 | DNA repair, bacterial RecFOR pathway | DNA processing |
| DNA recombination and repair protein RecO | 2 | DNA repair, bacterial RecFOR pathway | DNA processing |
| DNA recombination protein RmuC | 2 | DNA repair, bacterial | DNA processing |
| DNA repair helicase | 2 | Potential DNA repair cluster | DNA processing |
| DNA repair protein RadA | 2 | DNA repair, bacterial | DNA processing |
| DNA repair protein RecN | 2 | DNA repair, bacterial rRNA modification related cluster including tlyA | DNA processing |
| Endonuclease VIII | 4 | DNA repair base excision | DNA processing |
| Error-prone repair protein UmuD | 2 | DNA repair, bacterial UmuCD system | DNA processing |
| Excinuclease ABC subunit A | 4 | DNA repair, UvrABC system | DNA processing |
| Excinuclease ABC subunit A parlog of unknown function | 6 | DNA repair, UvrABC system | DNA processing |
| Excinuclease ABC subunit B | 4 | DNA repair, UvrABC system | DNA processing |
| Excinuclease ABC subunit C | 2 | BarA-UrvY(SirA) two component regulatory system DNA repair, uvrABC system | Regulation and Cell signaling<br>DNA processing |

**Table S1-3.**
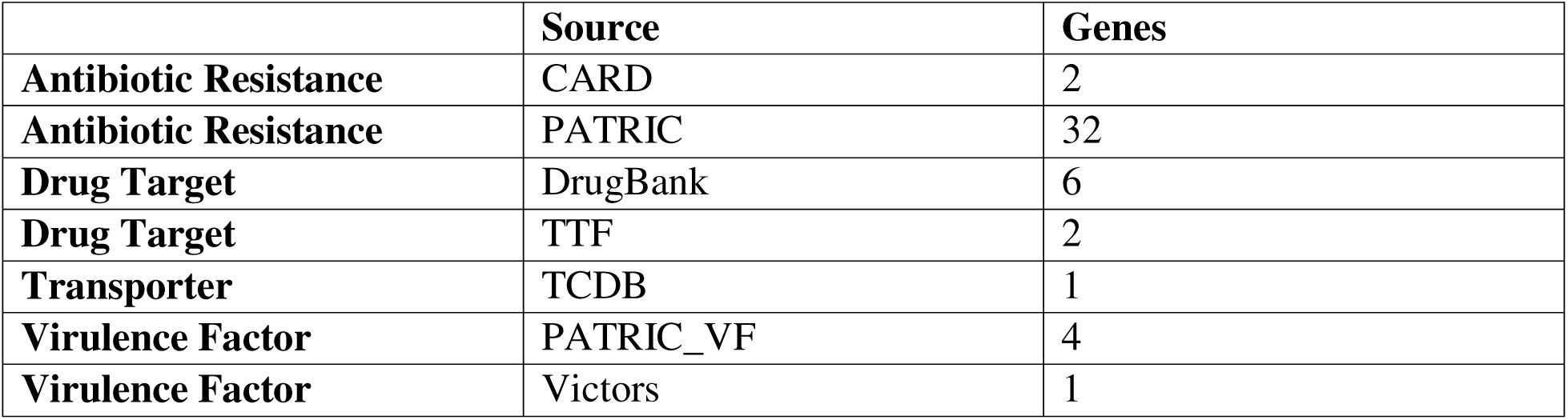
Number and types of specialized genes found in the MAG of *Arthrobacter oryzae*.

## Notes

### Competing Interest Statement

The authors have declared no competing interest.

